## Supplemental Figures 1-10 for "Machine Learning Discovers Numerous New Computational Principles Supporting Elementary Motion Detection"

**Supplementary figures**

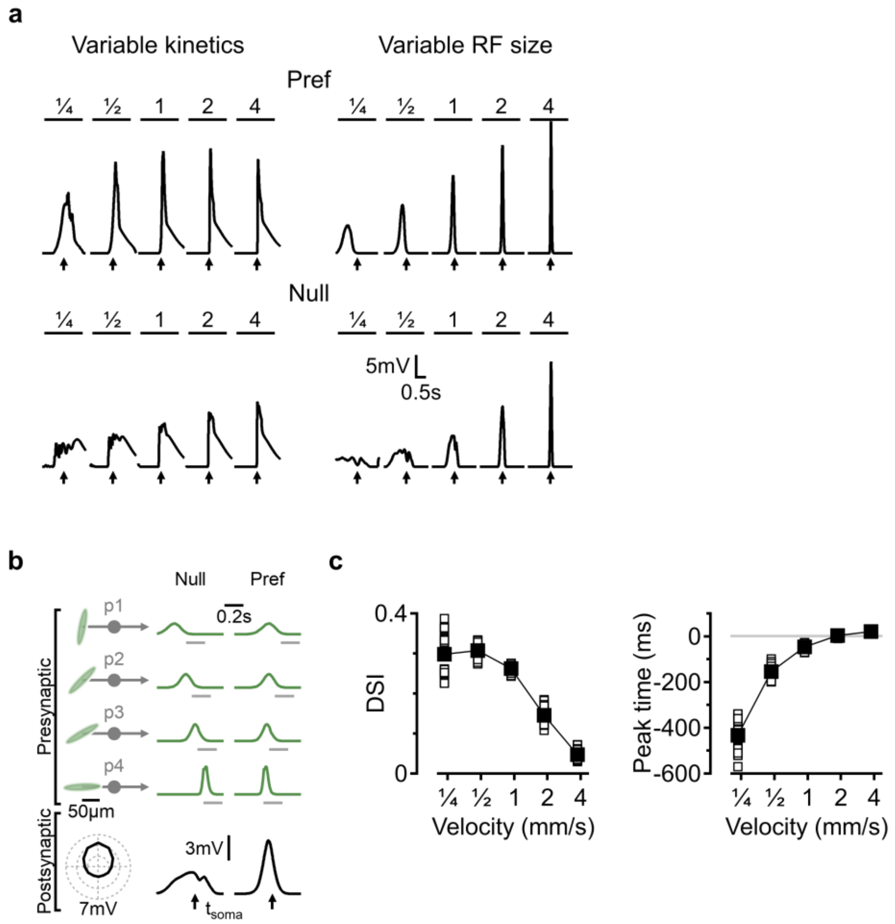


**Supplementary Figure 1: Comparison of direction selectivity models with varying center components of the presynaptic receptive fields. a**, Somatic voltage responses (top, preferred direction; bottom, null direction) across multiple velocities for the models in (**Fig. 1b, c**), showing earlier peak times in the spatial RF model. Arrows indicate the time when the stimulus reaches the position of the soma. **b**, Representative optimal solution to a full model with oriented presynaptic RFs. Top left, temporal RF parameters and the shape of the spatial RF were identical across all presynaptic cells. The free parameter in the model was the orientation of oval-shaped presynaptic RFs. Top right, normalized time courses of motion responses, with horizontal grey bars indicating the stimulation period. Bottom: membrane potentials and their directional tuning recorded in DSGC soma. $t_{soma}$ indicates the time point where the stimulus reached DSCG soma. **c**, Velocity tuning curves (left) and peak response times (right) in the preferred direction for 30 best-performing models evolved from unique initial parameters (open symbols), and population means (solid black symbols and lines). Negative values indicate responses occurring before the stimulus reaches the postsynaptic soma.

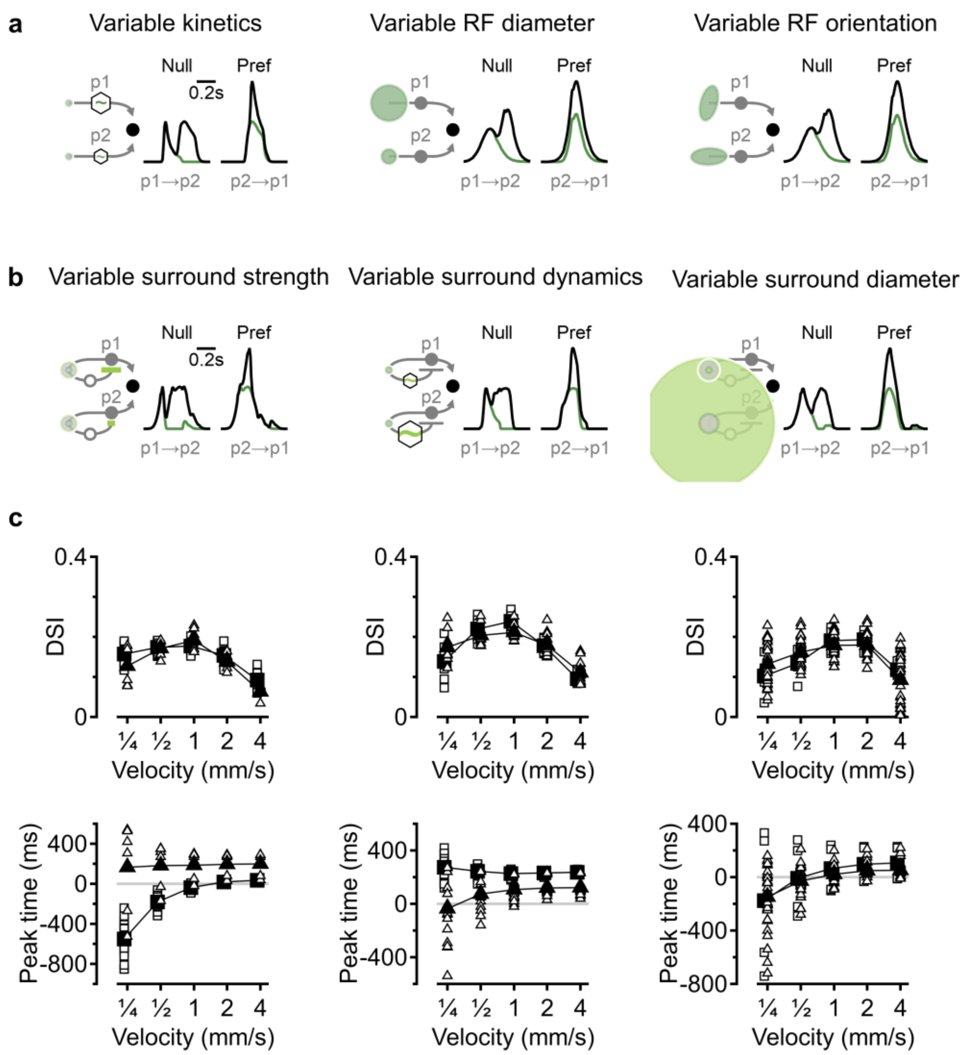


**Supplementary Figure 2: Models with temporal and spatial varying RF-centers and surrounds. a**, Representative responses in a minimal circuit consisting of two presynaptic neurons with varying RF center dynamics (left), size (center) or orientation (right) and a linear postsynaptic integrator. Black, detector response in the null/preferred direction of activation, green trace shows the response of the presynaptic neuron activated first in each sequence of stimulation. **b**, A representative solution of a minimal DS circuit model composed of presynaptic cells with varying strengths (left), kinetics (center), or size (right) of RF surrounds. RF centers and the spatiotemporal characteristics of the surround (but their weight in the full RF) were identical across the presynaptic populations. **c**, Velocity tuning curves (top) and peak response times (bottom) for the full models shown in (**Fig. 2a-c**). Triangles and squares correspond to distinct solution families calculated using hierarchical clustering of evolved model parameters. Representative examples from each cluster are depicted on the top and bottom panels in **Fig. 2a-c**, respectively.

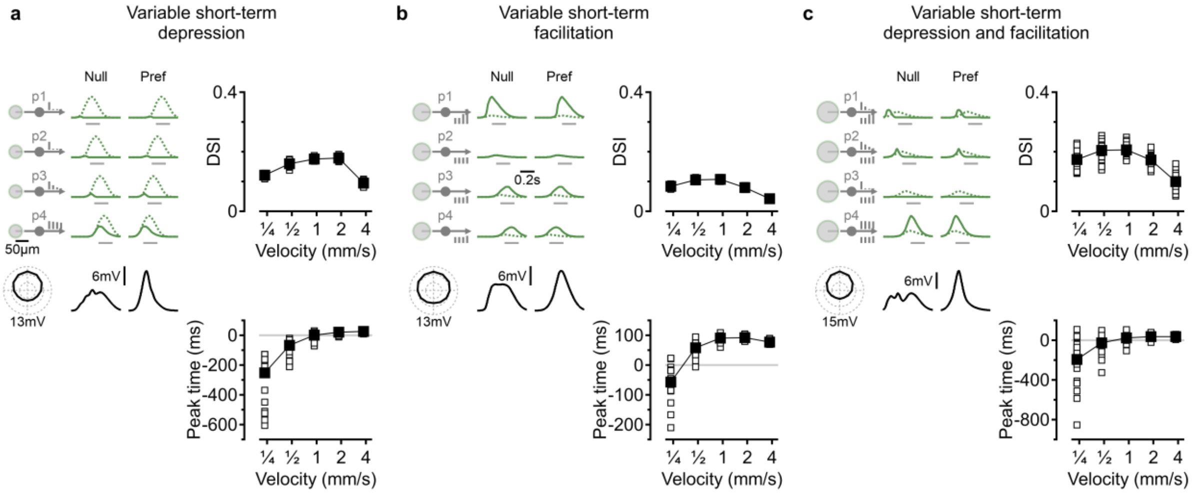


**Supplementary Figure 3: Models with short-term plasticity of the excitatory drive. a**, Left, example solution for DS circuit composed of presynaptic excitatory cells with identical RFs but different short-term synaptic depression levels. Dotted, RF activation prior to synaptic depression. The level of synaptic depression is depicted schematically for each input population. Right, velocity tuning (top) and peak response times (bottom). **b**, As in (**a**), but for varying short-term facilitation. **c**, As in (**a**), but for varying short-term facilitation and depression.

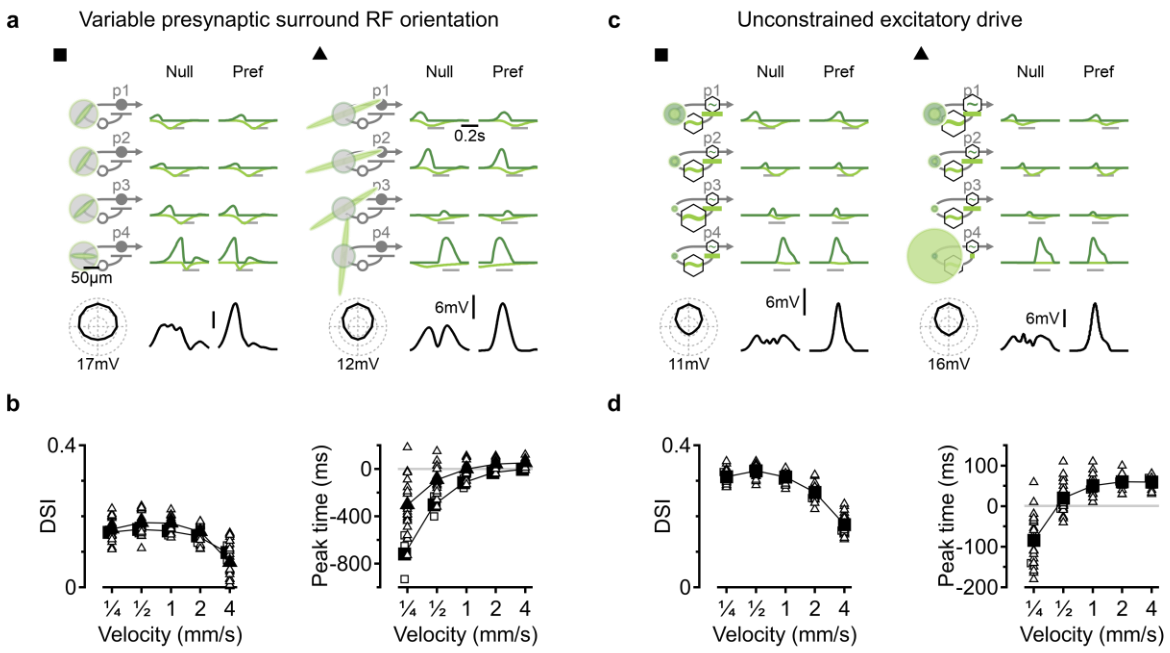


**Supplementary Figure 4: Direction selectivity in models with varying presynaptic RF surround orientation or unconstrained excitatory drive. a**, Two example solutions for the direction selectivity circuit composed of presynaptic cells with varying surround orientations. Spatiotemporal characteristics of the center, surround temporal dynamics and the spatial shape of the surround are identical between the inputs. **b**, Velocity tuning curves (left) and peak response times (right) of all optimal solutions (open symbols) and cluster means (black symbols and connecting lines); cluster assignment was calculated using hierarchical clustering. (**c**-**d**) As in (**a**-**b**) for selectivity circuit models where all spatiotemporal RF components were allowed to vary between the presynaptic populations.

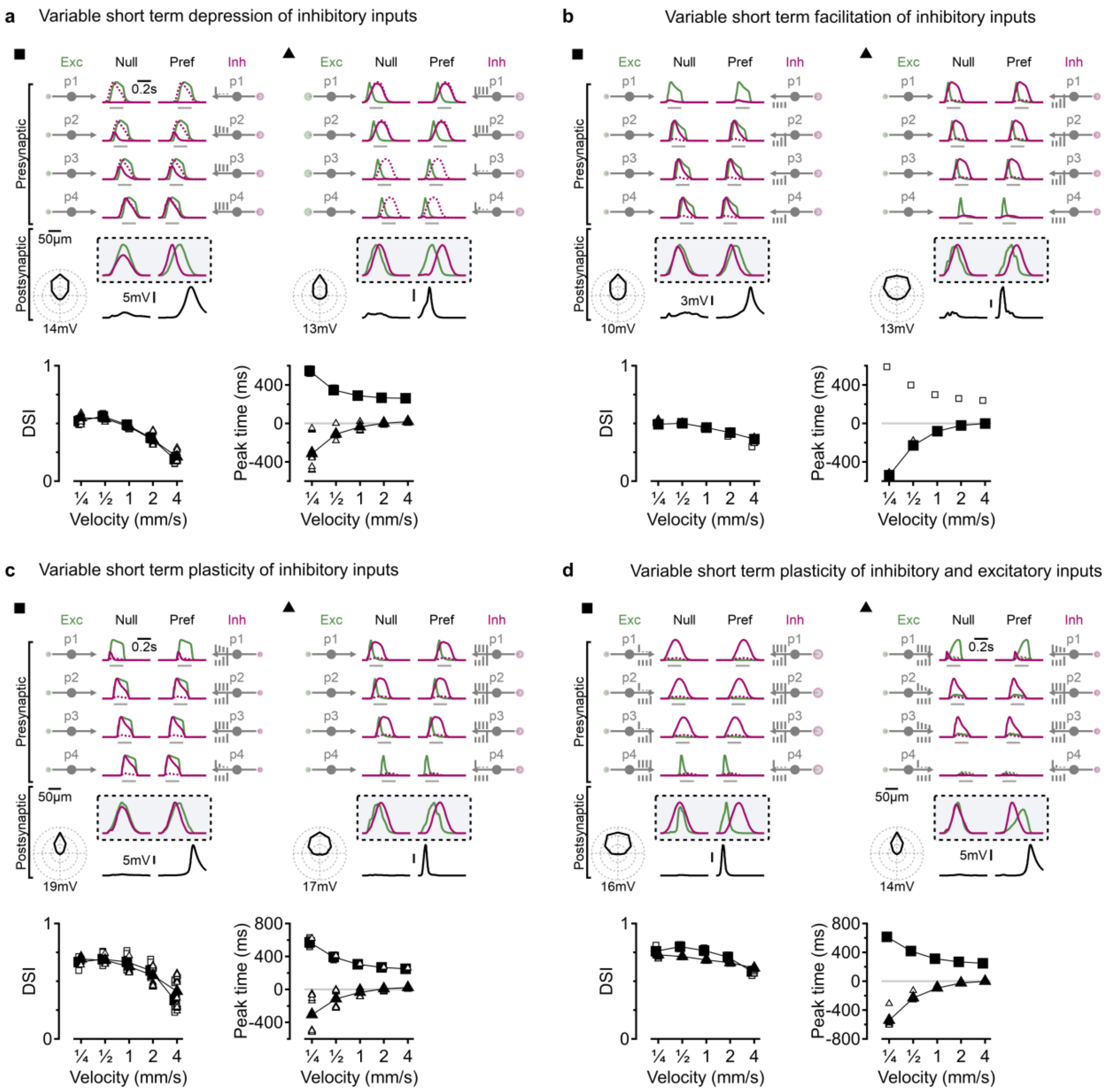


**Supplementary Figure 5: Models with short-term plasticity of the inhibitory drive. a**, Top, two representative solutions to DS circuits with presynaptic inhibitory cells having different short-term synaptic depression levels. Dotted, original RF activation. Solid curves, signals after filtering with depressing synapses. Green, left – excitatory inputs. Magenta, right – inhibitory inputs. Bottom, velocity tuning (bottom left) and peak response times (bottom right). Model responses were clustered by their evolved parameters, resulting in a separation between B&L-like (triangle) and anti-B&L (square) interactions. **b-d**, As in (**a**), but for varying short-term facilitation (**b**), short-term plasticity of the inhibitory population (**c**), and short-term plasticity implemented in all synaptic inputs (**d**).

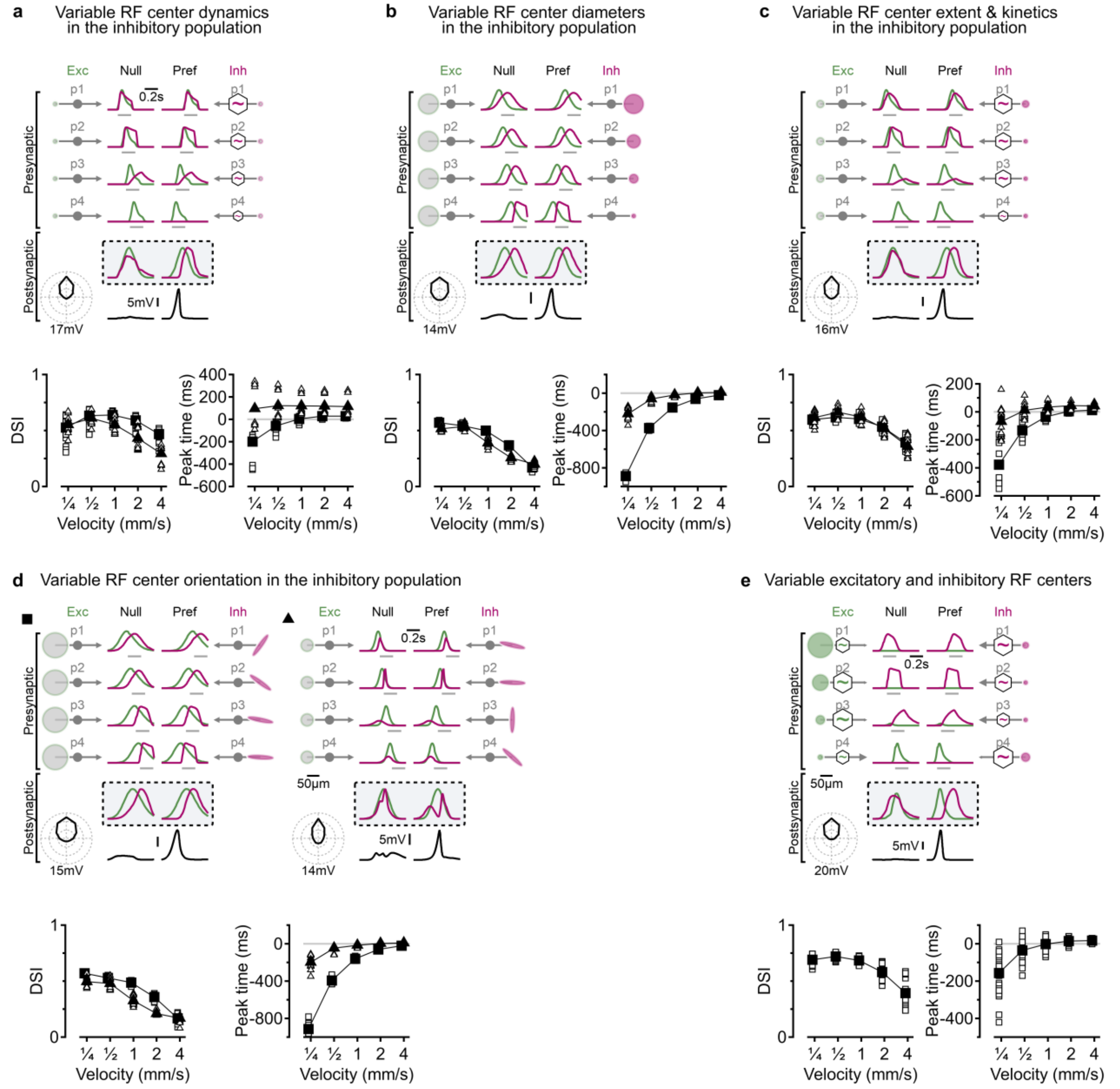


**Supplementary Figure 6: Models with varying spatiotemporal properties of inhibitory centers.** **a**, Top, a representative solution to direction selectivity circuit innervated by inhibitory cells with varying receptive field center dynamics. Green, left – excitatory inputs. Magenta, right – inhibitory inputs. Bottom, velocity tuning and peak response times for the preferred direction, separated into two clusters marked with squares and triangles based on the trained model parameters. **b-e**, As in (**a**), but for models with varying center-RF diameters of presynaptic inhibitory cells (**b**), variable RF size and kinetics (**c**), RF orientations (**d**), and variable RF size and kinetics of both excitatory and inhibitory drives (**e**).

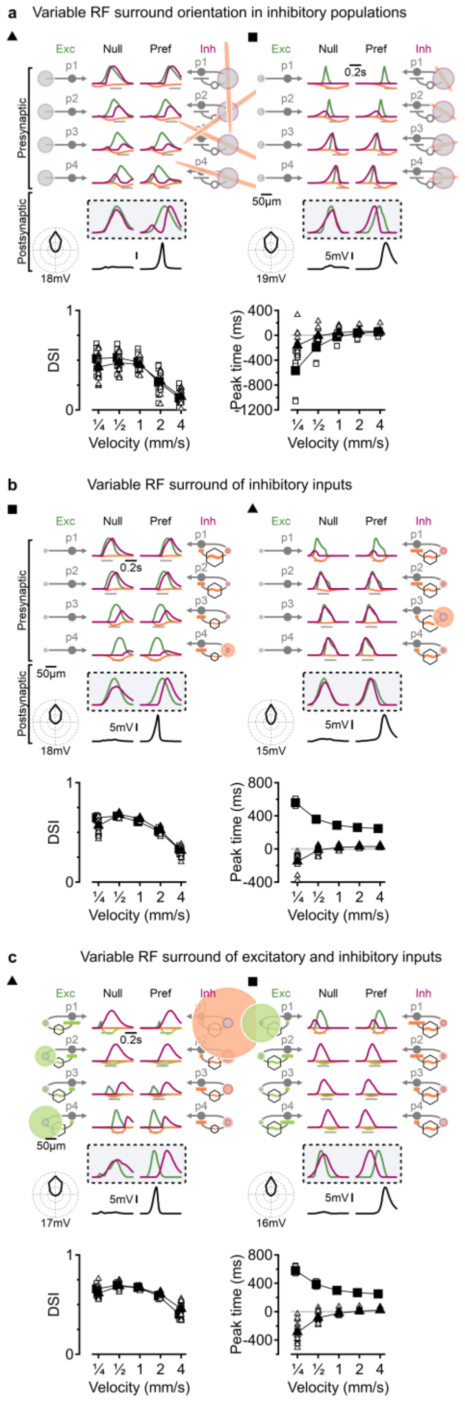


**Supplementary Figure 7: Solutions in models with varying spatiotemporal properties of surround receptive field characteristics of the presynaptic inhibitory cells. a**, Top, representative solutions with B&L-like (triangle) and anti-B&L (square) interactions observed in a direction selectivity circuit innervated by inhibitory cells with varying RF surround strengths. Bottom, velocity tuning (left), and peak response times (right) of the two solution types. **b**, As in (**a**) but for a variable description of inhibitory cells' surrounds. **c**, Models free to vary in surround parameters of excitatory and inhibitory cells.

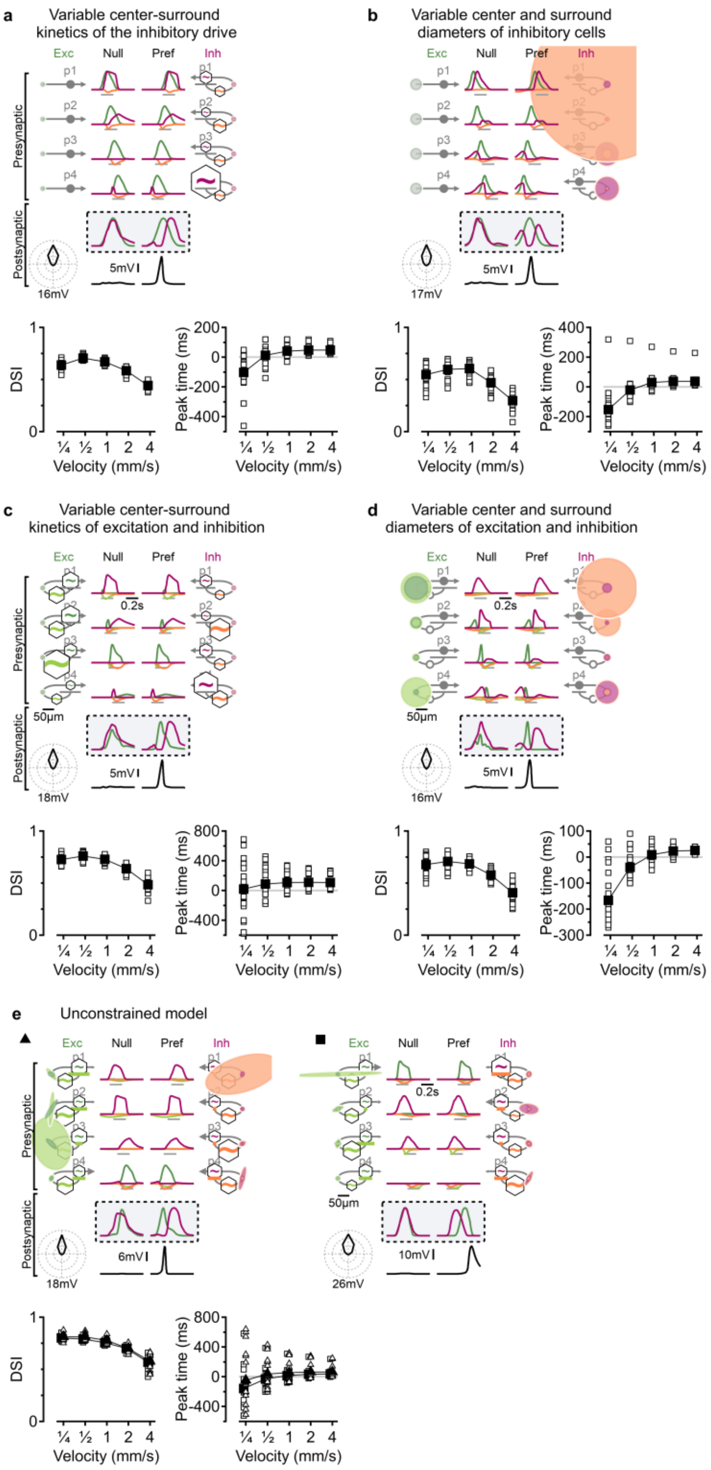


**Supplementary Figure 8: Solutions in models with multiple varying RF properties. a-b**, Top, a solution with unconstrained center-surround kinetics (**a**) and sizes (**b**) of the inhibitory cells. Bottom, velocity tuning (left), and peak response times (right). **c-d**, As in (**a-b**), but for varying excitation and inhibition. **e**, As in (**a**), but for models with unconstrained RF properties of excitatory and inhibitory populations exhibiting B&L (triangle) and anti-B&L (square) computational primitives.

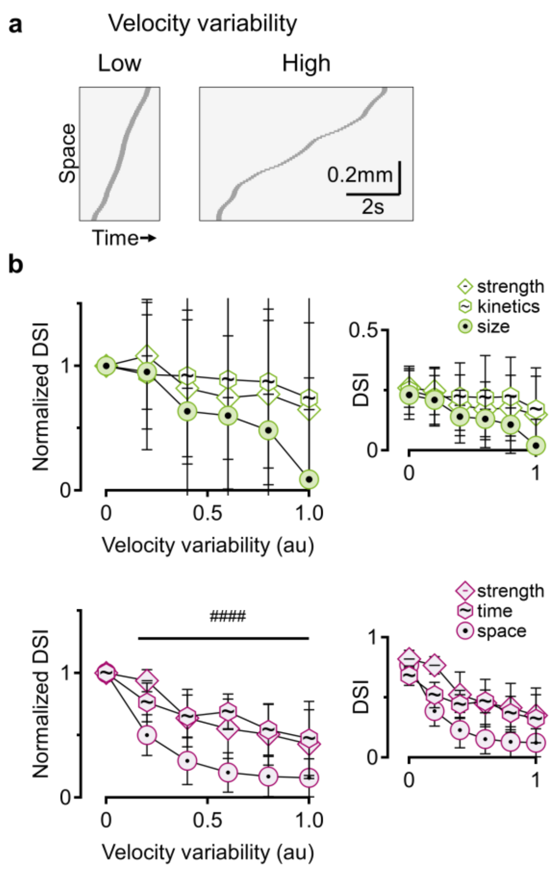


**Supplementary Figure 9. Differential resilience of surround DS solutions to stimulus variability. a**, Space-time plots as in **Fig. 9a**. **b-c**, Summary of directional discrimination as a function of stimulus variability for excitation-only models (**b**) and in the presence of inhibition (**c**). DSI values were normalized to the no-noise baseline (inset, absolute DSI values). ####p < 0.0001; significant difference between noise-dependence slopes of variable inhibitory RF surround extents and other inhibitory configurations (ANOVA on linear regression fits, n = 100). Error bars: SD.

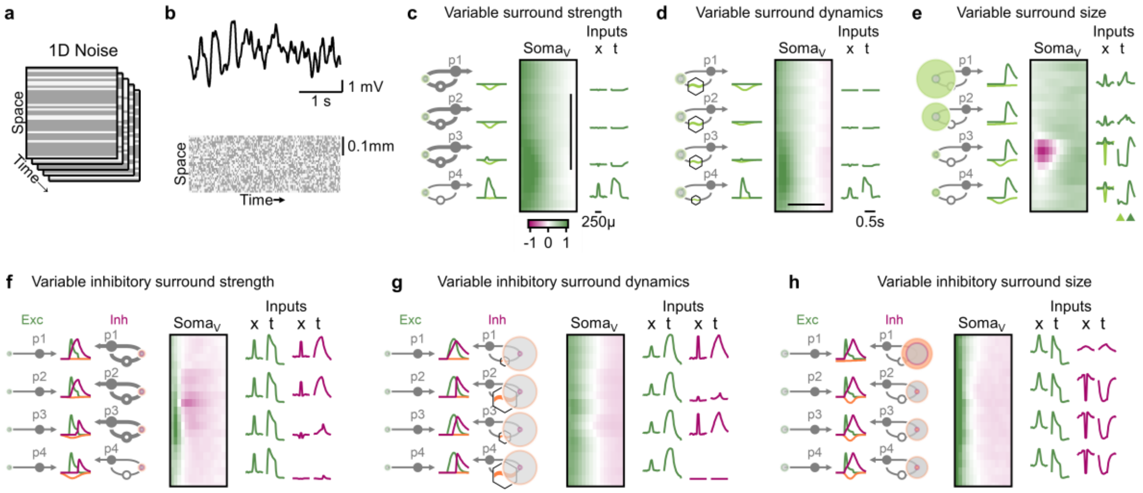


**Supplementary Figure 10. 1D noise stimulation can not reliably resolve the architecture of DS circuits that depend on surround RF components.** As in **Figure 10**, receptive field structures were assessed using oriented bars aligned with the preferred direction of motion. **a**, In each frame, individual bars were randomly assigned black or white contrast values to create a 1D bar noise stimulus. **b**, Representative membrane potential trace recorded from the DSGC soma (top) in response to 1D bar noise stimulation (bottom) in (**a**) model where motion selectivity arises from differences in surround strengths. **c**, Left: Ground-truth spatiotemporal profiles of four excitatory presynaptic populations from an evolved model with variable surround strengths. Temporal responses were derived from a full-field flash stimulus. Center: Space-time plot of the average postsynaptic response in DSGC following the appearance of a white bar at each spatial location. The vertical axis indicates spatial position; the horizontal axis shows time after stimulus onset. Color coding represents depolarization (green) and hyperpolarization (red). Scale bar: 250 µm, centered over the DSGC's RF. Right: Inferred RF properties of the presynaptic populations based on their responses to the 1D noise stimulus. Presynaptic analysis captures some circuit organization principles but does not reveal the role of surround. **d-e**, As in (**c**), but for models in which motion selectivity arises from differences in surround RF dynamics (**d**) size (**e**). Colored triangles in (**e**) indicate two different time points for which spatial RF was analyzed. **f-h,** As in (**c**), for models in which DS is mediated by differences in inhibitory surround strength (**f**), dynamics (**g**), or size (**h**).
